# Short-term consumption of ultra-processed semi-synthetic diets impairs the sense of smell and brain metabolism in mice

**DOI:** 10.1101/2023.08.31.555480

**Authors:** Melanie Makhlouf, Débora G. Souza, Smija Kurian, Bruna Bellaver, Hillary Ellis, Akihito Kuboki, Asma Al-Naama, Reem Hasnah, Gianina Teribele Venturin, Jaderson Costa da Costa, Neethu Venugopal, Diogo Manoel, Johannes Reisert, Michael G. Tordoff, Eduardo R. Zimmer, Luis R. Saraiva

## Abstract

The prevalence of highly-palatable, ultra-processed food in our modern diet has exacerbated obesity rates and contributed to a global health crisis. While accumulating evidence suggests that chronic consumption of ultra-processed semi-synthetic food is detrimental to sensory and neural physiology, it is unclear whether its short-term intake has adverse effects. Here, we assessed how short-term consumption (<2 months) of three ultra-processed diets (one grain-based diet, and two semi-synthetic) influence olfaction and brain metabolism in mice. Our results demonstrate that short-term consumption of semi-synthetic diets, regardless of macronutrient composition, adversely affect odor-guided behaviors, physiological responses to odorants, transcriptional profiles in the olfactory mucosa and brain regions, and brain glucose metabolism and mitochondrial respiration. These findings reveal that even short periods of ultra-processed semi-synthetic food consumption are sufficient to cause early olfactory and brain abnormalities, which has the potential to alter food choices and influence the risk of developing metabolic disease.

## INTRODUCTION

The evolutionary history of modern humans is riddled with seismic shifts in their patterns of food consumption and physical activity [1, 2]. Presently, most humans in Western societies live sedentary lives in a fast-paced world and are constantly bombarded with myriad sensory stimuli, which encourage over-consumption of a wide variety of energy-dense and ultra-processed unhealthy foods [3-5]. This over-consumption, when sustained for long periods, leads to the dysregulation of the appetite system, weight gain, and the development of a wide range of noncommunicable diseases – ranging from metabolic disorders to cancer, autoimmune diseases, mental and brain health problems [2, 6-9].

Sensory inputs, such as food odors, interact with internal homeostatic and hedonic signals in the brain to regulate food retrieval and consummatory behaviors [4, 10-13]. In appetite regulation, olfaction is perhaps the most evocative of all five senses. Indeed, several lines of evidence support the essential role of smell in food detection and then selection, which determines dietary quality, caloric intake, and metabolic health [14-16]. Additionally, olfactory dysfunction (e.g., hyposmia) further stimulates the ingestion of nutrient-poor and energy-dense diets, leading to an unhealthy vicious cycle [17, 18]. In turn, the long-term consumption of energy-dense diets (e.g., high-fat or high-fructose) can negatively impact olfactory physiology, odor-guided behaviors, and lead to altered brain function in regions involved in olfactory processing [4, 10, 11, 19-25].

Considering the major shifts in eating culture, the prevalence of cheap and readily-available fast food, and resulting obesity epidemic over the last five decades, it is crucial to map and better understand the effects of not just long-term but also short-term consumption of semi-synthetic diets on health and disease. Interestingly, a recent study showed that mice fed a semi-synthetic diet have a lower ability to defend against influenza when compared to animals fed a standard mouse chow diet with similar macronutrient composition [26]. Moreover, recent studies of mice have shown that consuming semi-synthetic high-fat diets for only a few days can lead to metabolic changes (e.g., ketogenesis, hepatic steatosis, insulin resistance), and alter the transcriptional profile of the intestine [27-31]. Despite these recent advances in the field, how the sense of smell and brain function are affected by the short-term consumption of semi-synthetic diets remains largely unknown. To address this question, we used mice to investigate the effects of the short-term consumption (<8 weeks) [32, 33] of one grain-based processed diet or two semi-synthetic diets on odor-guided behaviors, olfactory physiology, the transcriptional profiles in the olfactory mucosa and various brain regions, as well as brain glucose metabolism and mitochondrial respiration.

## RESULTS

### Short-term consumption of semi-synthetic diets severely impacts odor-guided behaviors

Long-term unhealthy dietary interventions can lead to impairments in memory, and emotional or cognitive behaviors (such as reward-, social- and stress-related behaviors), even independently of obesity [9, 21, 24, 34-37]. The long-term consumption of high-fat, high-fructose, or Western diets can also lead to defects in specific olfactory-related behaviors, such as odor discrimination, odor-related learning, and memory [21, 22, 25, 37, 38]. However, whether the short-term consumption of semi-synthetic diets can affect the sense of smell remains unknown. To address this question, we randomly allocated C57Bl/6J male mice to one of the three ultra-processed diets that differ in ingredient origin and macronutrient composition: a grain-based “normal” chow diet (NCD), a semi-synthetic control diet (SCD), or a semi-synthetic high-fat diet (HFD) (see STAR Methods, Figure 1A, Data File S1). Mice fed the HFD weigh significantly more than mice on the NCD or the SCD (Figure S1). After being on their designated diet for three weeks, we performed olfactory preference behavioral tests against an odorless control (water, H2O) and nine odorants (hexanethiol, HXT; linalool, LIN; hexanol, HXO; isoamylamine, IAA; vanillin, VAN; indole, IND; (+)-menthone, +MNT; trimethylamine, TMA; ethyl butyrate, EBT) that have been shown to elicit a range of behavioral outputs in mice [39, 40]. To avoid habituation, each mouse was used only once and against one odorant (n = 7-12 mice per odorant) and scored on three different behavioral parameters surrogate for odor “valence” (OIT, olfactory investigation time), stress (RAS, risk assessment), and exploration (VEL, velocity) (Figure 1B). This dataset totaled 722 data points corresponding to individual-odorant-behavior triads (Figures 1C-E and S1A, Data S1), for which we calculated the average value for each of the 30 odorant-behavior pairs for each tested diet. Surprisingly, the patterns of significant increases and decreases compared to H2O (p<0.05; one-way ANOVA; BKY multiple-comparisons correction) across the different behaviors differ across the three different diets, with NCD displaying the highest number of differences (10), followed by SCD (4) and HFD (2) (Figure 1F). Only 2 of the 16 significant odorant-induced behavioral changes were shared between at least two diets: the significant decrease of OIT for HXT (in NCD and SCD) and the significant increase of RAS for LIN (in NCD and HFD). Next, we compared the same odorant-behavior pairs between diets (Figure 1G), and found unique behavioral differences amongst pairwise diet comparisons. The highest number of significant behavioral differences (13) was observed in the SCD vs. NCD, followed by 8 in HFD vs. NCD, and 4 in HFD vs. SCD. In both comparisons vs. NCD, most OIT and VEL differences are significantly higher in NCD, but all RAS differences are significantly higher in SCD and HFD. In HFD vs. SCD, the only OIT difference is significantly higher in HFD, and the differences in RAS and VEL are all significantly higher in SCD. Moreover, each diet displayed unique patterns of significant changes among the three behaviors tested. Together, these results indicate that short-term dietary interventions can substantially impact odor-guided behaviors in mice, with semi-synthetic diets strongly impacting their standard olfactory preferences, and mildly increasing their stress-related behavior

**Fig 1.**
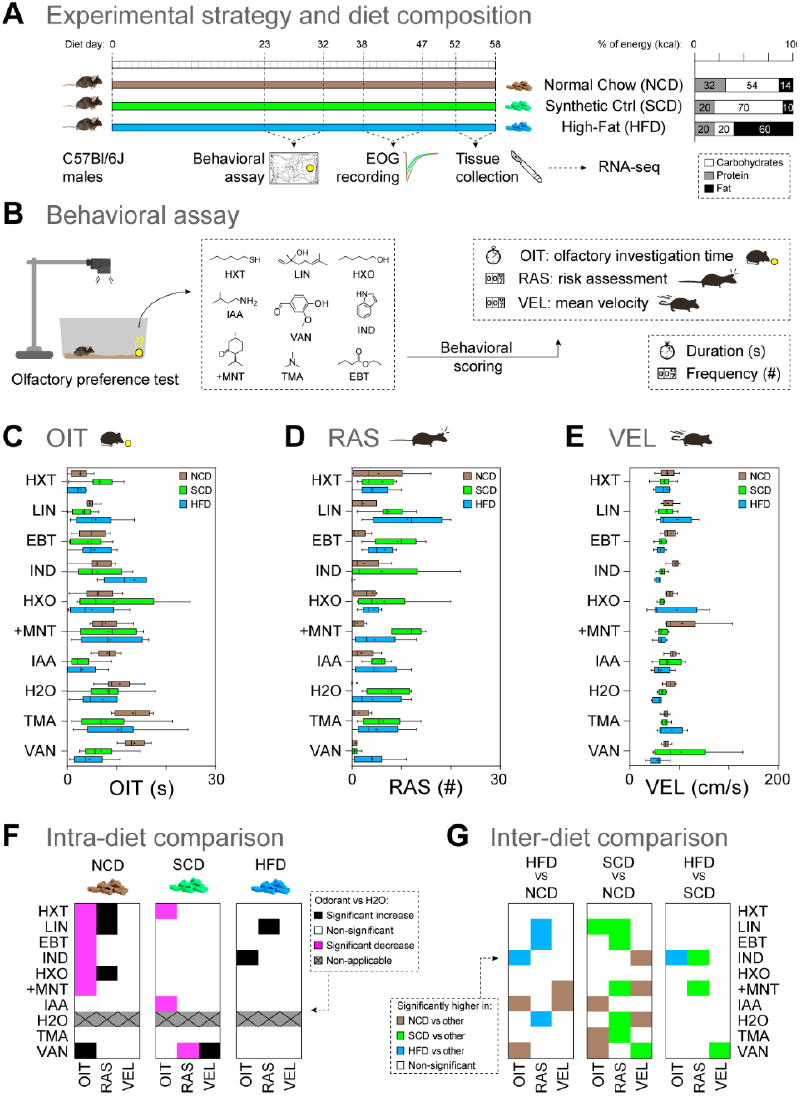
Effects of short-term consumption ofdiets differing in macronutrient origin and composition on odor-guided behaviors.

(A) Schematic view of the experimental design for the first part of our study, comprising three different diets: a grain-based normal chow diet (NCD, brown bar), the semi-synthetic control diet (SCD, green bar), and the semi-synthetic high-fat diet (HFD, blue bar). The macronutrient composition (%) for each diet is depicted on the right (carbohydrates in white, protein in grey, and fat in black). (B) Behavioral assays: animals were exposed to nine odorants, and videos were recorded and scored for three distinct behavioral parameters: olfactory investigation time (OIT), risk assessment (RAS), and velocity (VEL). Odorants used: (+)-menthone (+MNT), ethyl butyrate (EBT), hexanethiol (HXT), hexanol (HXO), indole (IND), isoamylamine (IAA), linalool (LIN), trimethylamine (TMA), vanillin (VAN). (C-E) Boxplots summarizing the behavioral parameter scores for the nine tested odorants. OIT scores are shown in seconds (C), RAS scores in frequencies (D), and VEL scores in distance per second (E).(F) A graphical display in the form of a heatmap summarizing the statistical significance results of intra-diet comparisons. Behaviors showing significant (one-way ANOVA, BKY multiple comparisons correction, n = 7-12 per odorant) increases, decreases and non-significant responses compared to H2O (no odor control) are indicated in black, magenta, and white squares, respectively. Non-applicable comparisons are in grey. (G) A graphical display in the form of a heatmap summarizing the statistical significance of inter-diet comparisons for each odorant-behavioral parameter dyad. Behaviors with significantly higher values (one-way ANOVA, BKY multiple comparisons correction, n = 7-12 per odorant) in NCD, SCD or HFD diets are indicated in brown, green or blue, respectively. Non-significant responses are indicated in white.

### The short-term consumption of semi-synthetic diets impairs odorant detection in the nose

Internal metabolic signals can directly modulate the response of olfactory sensory neurons (OSNs) to odorants [20, 41]. Thus, the behavioral deficits observed in animals fed the semi-synthetic SCD and HFD could be due, at least in part, to changes in the peripheral detection of odorants in the neuroepithelium of the mouse olfactory mucosa. To test this hypothesis, we performed air-phased electro-olfactogram (EOG) recordings in response to the nine odorants used in the behavioral assays, two control odorants (pentyl acetate, PAC; dimethyl sulfoxide, DMSO) (Figure 2A). The amplitudes of the odorant responses varied between the three tested diets, with mice on the NCD displaying significantly larger EOG amplitudes to most odorants compared to SCD or HFD (p<0.05; two-way ANOVA, odorants: F(13,266) = 108.6, p < 0.0001; Diet: F(2,266) = 49.16, p < 0.0001; Interaction: F(26,266) = 1.514, p = 0.0564; BKY multiple-comparisons correction; Figure 2B).

**Fig 2.**
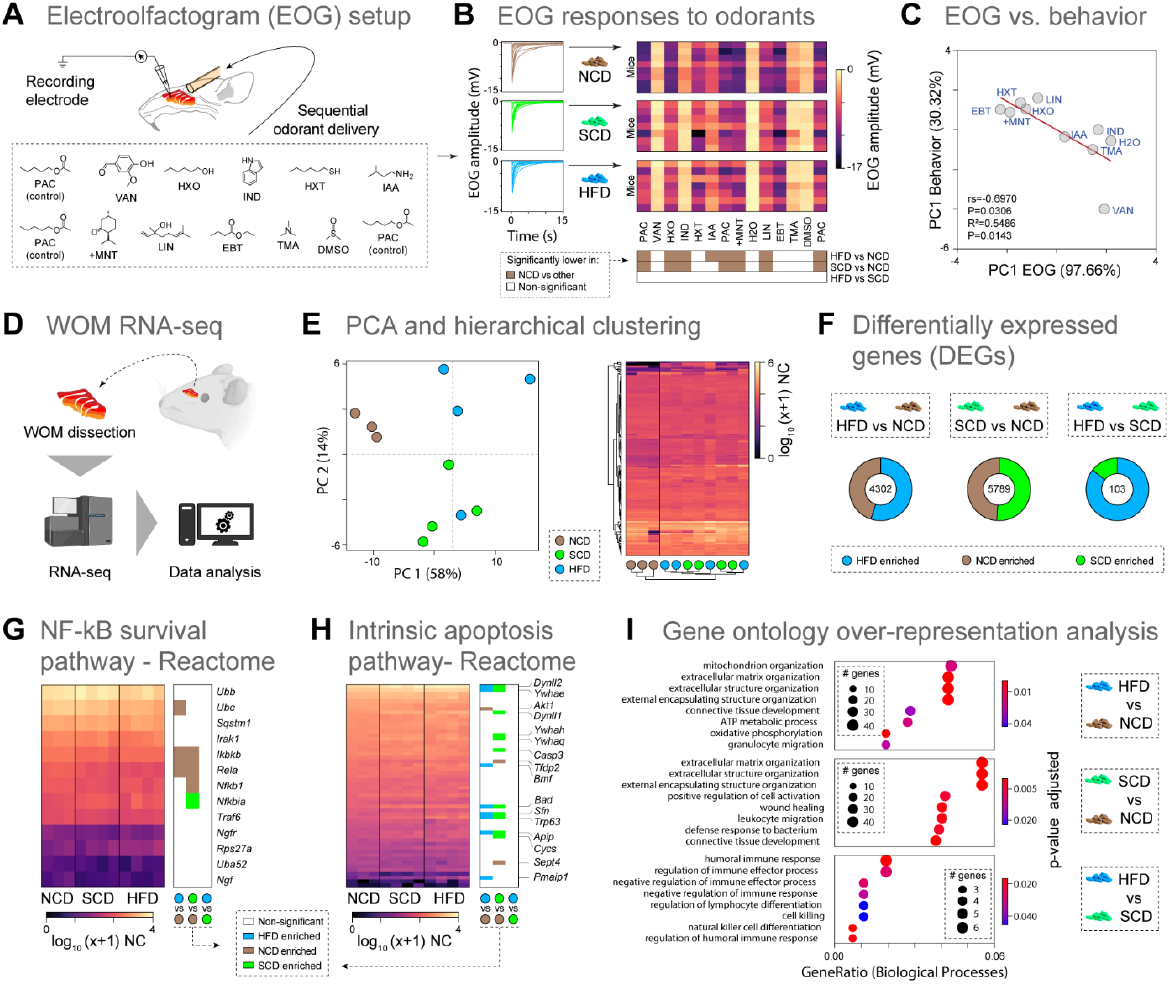
Semi-synthetic diets affect odorant detection and induce transcriptional dysregulations in the mouse nose.

(A) Schematic of the air-phased electroolfactogram (EOG) recording experiments performed with 11 odorants on mice exposed to the three test diets (NCD, SCD, and HFD). (B) Left panel: Odorant-evoked EOG response curves to odorants. Upper right panel: Heatmap showing the odorant evoked EOG amplitude values (mV) per mouse. Lower right panel: Heatmap summarizing the statistical significance of the comparison of peak amplitudes in EOG recordings in response to each odorant between diets. EOG amplitudes showing significantly lower values (p<0.05; two-way ANOVA; BKY multiple comparisons correction, n = 7-8 per odorant) in NCD, SCD or HFD diets are indicated in brown, green or blue, respectively. Non-significant responses are indicated in white. (C) Spearman correlation (rs) and linear regression (R2) between principal component 1 (PC1) of the EOG responses and PC1 of the odor-guided behaviors. (D) Schematic view of the RNA-seq experimental strategy for the whole olfactory mucosa (WOM) samples (n=3-4 mice per group). (E) PCA and hierarchical clustering of the union of the top 1000 most expressed genes across all WOM samples. Heatmap RNA-seq expression values are represented on a log10(x+1) scale of normalized counts (NC). Each dot represents an animal. (F) Pie chart displaying the total number and relative proportion of differentially expressed genes (DEGs; p-adj<0.05) for each pairwise diet comparison in the WOM. Genes enriched in mice fed on HFD, NCD or SCD are represented in blue, brown, or green, respectively. (G-H) Heatmap (left panel) showing the normalized gene expression levels of Reactome “NF-κB is activated and signals survival” pathway (G) or the Reactome “intrinsic apoptosis” pathway genes (H). Right panels: a graphical display in the form of a heatmap summarizing the statistical significance of inter-diet comparisons. DEGs enriched in mice fed on HFD, NCD, or SCD are represented in blue, brown, or green, respectively. (I) Dot plot representing selected significant terms from Gene Ontology (GO) over-representation analysis done on DEGs (|log2(FC)|>1, p-adj<0.05) from differential expression analysis between diets. Enriched terms are displayed by gene ratio (number of genes related to a GO term/total number of significant genes).The size of the dot reflects the number of genes enriched within the associated GO term, and its color reflects the p-adjusted value for each term.

Interestingly, there were no significant differences in EOG amplitudes between SCD and HFD, similar to our behavioral results. Together, our data suggest that deficits in peripheral detection of odorants is a result of the synthetic nature of the food rather than its macronutrient content.

Because we performed behavioral and physiological experiments in the same mice, we next asked whether the dietary-induced changes in odorant detection in the nose are associated with the corresponding odor-guided behaviors. We performed principal component analysis (PCA) on the data matrixes from the EOG responses and the odor-guided behaviors of the nine tested odorants and water (see STAR Methods, Figures S2A,B). Strikingly, we observed a strong correlation (rs=-0.69, p=0.03) between the PC1 (97.66% of the variance) of the EOG responses and the PC1 (30.32% of the variance) of the odor-guided behaviors (Figure 2C). The linear regression analysis (R2=0.55, p=0.01,) indicates that the decrease in responsiveness of OSNs to odorants, caused by consuming semi-synthetic diets, accounts for ∼55% in the variation of odor-guided behaviors. Overall, these findings demonstrate that short-term consumption of semi-synthetic food impairs olfactory processing and related behaviors. However, the impairment of olfactory function in the periphery does not fully account for its impact on odor-guided behaviors, suggesting the presence of other contributing mechanisms, likely happening in the brain.

### Semi-synthetic diets induce transcriptional dysregulations associated with apoptosis and inflammatory responses in the mouse nose

To better understand the mechanisms underlying the physiological dysregulation of odor perception in the nose, we performed RNA-sequencing (RNA-seq) in the whole olfactory mucosa (WOM) of the mice fed the NCD, SCD, or HFD (Figure 2D). PCA and hierarchical clustering (HC) analyses of the transcriptomic data segregated the samples according to the diets and into two groups: one included all samples from animals fed the grain-based NCD, and the other comprised the samples from animals fed the two semi-synthetic SCD and HFD (Figure 2E) – congruent with the fact that these two diets similarly impair olfaction (Figures 1 and 2). To gain more insights into the diet-induced transcriptomic changes, we performed differential expression analysis for all possible dietary pairwise comparisons. The comparisons between NCD and the semi-synthetic HFD and SCD yielded 4302 and 5789 differentially expressed genes (DEGs), respectively, but the comparison between SCD and HFD only yielded 103 DEGs (Figure 2F). Next, we focused on the canonical markers of the different cell types populating the WOM. We found subtle enrichment in the expression of markers for mature and immature OSNs (mOSNs and iOSNs, respectively) in WOMs from mice fed an NCD, and in the expression of horizontal basal cells (HBCs) markers in WOMs from mice fed a SCD or HFD (Figure S2C).

The changes in the expression of cell markers observed above could reflect differences in the abundance of those specific cell types caused by diet-induced dysregulation of neuroregenerative, inflammatory, or apoptotic processes [23, 25, 42]. To test this hypothesis, we asked whether genes involved in eliciting neuroregeneration or apoptosis are among the DEGs identified above. First, we looked at genes integrating the NF-κB-mediated cell survival pathway, which promotes neuroregeneration induced by acute inflammation [43]. We found that 5/13 are differentially expressed between NCD and HFD or SCD, with the majority (∼80%) being enriched in NCD compared to SCD or HFD (Figure 2G). Second, we analyzed the genes composing the intrinsic apoptotic pathway and found that 16/54 are differentially expressed between NCD and HFD or SCD, with the majority (∼82%) being enriched in SCD or HFD compared to NCD (Figure 2H). Together these results suggest that the WOMs of animals fed semi-synthetic HFD and SCD have higher levels of apoptosis than the WOMs of animals fed the NCD.

To gain deeper and unbiased insights into the functions of all diet-induced DEGs, we performed gene ontology (GO) over-representation analysis (see STAR Methods). We obtained a total of 18, 74, and 12 GO Biological Process categories enriched for the HFD vs. NCD, SCD vs. NCD, and HFD vs. SCD, respectively (Data File S2). Three of the top five GO-enriched categories for the HFD vs. NCD and SCD vs. NCD comparisons are related to the extracellular matrix and structure organization, with most genes composing these three categories being enriched in NCD compared to SCD or HFD (Figure 2I, Data File S2). Interestingly, lower expression levels of extracellular matrix genes are associated with lower numbers of collagen-expressing mesenchymal cells and age-dependent impairment of olfaction [42], suggesting that the WOMs of animals fed an NCD may have higher regenerative potential than the WOMs of animals fed semi-synthetic diets. In contrast, genes displaying enriched expression in SCD or HFD compared to NCD belong to GO categories involved in ATP metabolic processes, oxidative phosphorylation, wound healing, and immune function (Figure 2I, Data File S2); suggesting that the WOMs of animals fed semi-synthetic diets have higher levels of inflammation and tissue damage. Lastly, most categories enriched between HFC and SCD are related to the immune system and composed of genes showing enriched expression in HFD compared to SCD (Figure 2I, Data File S2); thus, suggesting a higher level of inflammatory processes in animals fed HFD, which then trigger enhanced immune system responses. Together, these results indicate that the short-term consumption of semi-synthetic diets negatively affects cellular processes regulating neuroregeneration, inflammation, and cell death, consistent with changes observed in aged olfactory epithelia [42-44], and with previous studies performing long-term dietary interventions [22, 23, 25].

### Short-term consumption of semi-synthetic diets differentially affects the transcriptional profiles of various brain regions

Internal metabolic signals can also modulate olfaction by acting on the central nervous system (CNS) [10, 20, 41]. Furthermore, consuming different macronutrients in the diet can modulate gene expression patterns and thereby affect brain physiology [37, 45, 46]. Our results above indicate that the decrease in responsiveness of OSNs to odorants, caused by consuming semi-synthetic diets, only accounts for half of the variation of odor-guided behaviors (Figure 2C). These data suggest the presence of additional modulatory mechanisms in the brain, which are likely regulated by diet-induced changes in gene expression and contribute to the adverse effects of ynthetic-ingredient-based diets on odor-guided behaviors.

To test this hypothesis, we performed RNA-seq in four broad brain regions — olfactory bulb (OB), brain (BRN, consisting of the cortical and subcortical regions), cerebellum (CBL), and brainstem (BST, which consists of the mesencephalon, pons, and myelencephalon) — of mice fed the NCD, SCD, or HFD (Figure 3A, see STAR Methods). As expected, PCA and hierarchical clustering analysis of the transcriptomic data segregated the samples first according to brain regions and second according to the diet (Figures 3B and S3A).

**Fig 3.**
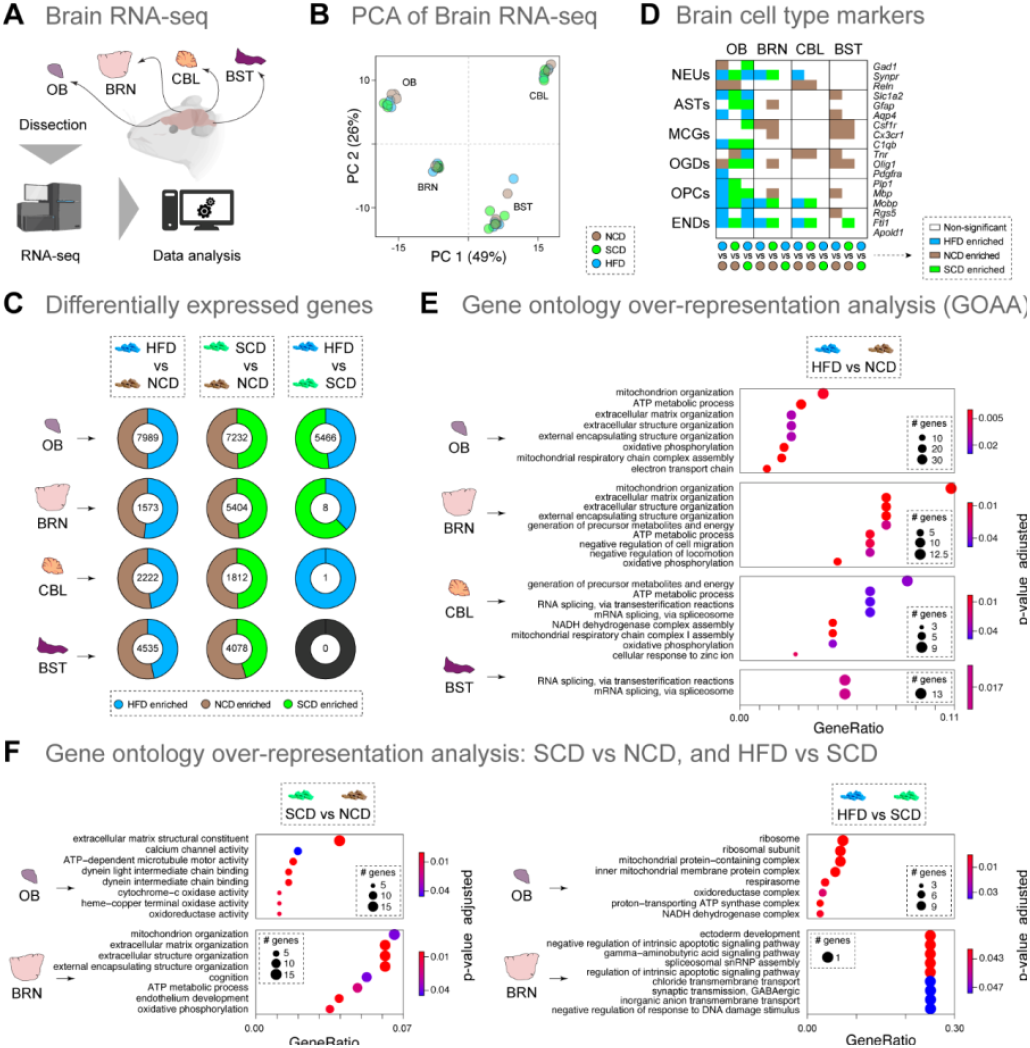
Semi-synthetic diets differentially affect the transcriptome of various brain regions. (A) Schematic view of the RNA-seq experimental strategy for the four broad brain regions: olfactory bulb (OB), brain (BRN, consisting of the cortical and subcortical regions), cerebellum (CBL), and brainstem (BST, which consists of the mesencephalon, pons, and myelencephalon) of mice fed different diets (NCD, SCD, HFD n=3-4 mice per group). (B) Principal-component analysis (PCA) of the union of the top 1000 most expressed genes across all brain regions samples. (C) Pie chart showing the total number and relative proportion of differentially expressed genes (DEGs; p-adj<0.05) for each pairwise comparison between diets within the different brain regions. Genes enriched in mice fed on HFD, NCD, or SCD are represented in blue, brown, or green, respectively. (D) Heatmap highlighting differentially expressed genes between diets and within different brain regions for known markers of brain cell types (NEUs, neurons; ASTs, astrocytes; MCGs, microglial cells; OGDs, oligodendrocytes; OPCs, oligodendrocyte progenitor cells; ENDs, endothelial cells). DEGs enriched in mice fed on HFD, NCD, or SCD are represented in blue, brown, or green, respectively. Genes with no significant expression changes are represented in white. (E) Dot plot of selected significant Gene Ontology (GO) terms performed on DEGs (|log2(FC)|>1, p-adj<0.05) from HFD vs. NCD diet comparison in OB, BRN, CBL or BST regions. (F) Dot plot of selected significant Gene Ontology terms obtained by gene over-representation analysis done on DEGs (|log2 (FC)|>1, p-adj<0.05) from SCD vs. NCD (left panels) or HFD vs. SCD (right panels) diet comparisons in OB and BRN regions. Enriched terms are displayed by gene ratio (number of genes related to a GO term/total number of significant genes). The size of the dot reflects the number of genes enriched within the associated GO term, and its color reflects the p-adjusted value for each term.

To gain more insights into the diet-induced transcriptomic changes, we performed differential expression analysis for all possible dietary pairwise comparisons. Across all brain regions, the dietary comparisons yielding the highest number of DEGs were between NCD and the semi-synthetic SCD and HFD, ranging from 1573 to 7989 DEGs (Figure 3C). The comparison between SCD and HFD yielded 5466 DEGs for the OB, 8 for BRN, 1 for CBL, and none for BST. Interestingly, of the four brain regions analyzed, the OB registered the highest number of DEGs across all the dietary comparisons (20,687), followed by the BST (8,613), BRN (6,985), and CBL (4,035). Next, we focused on the canonical markers of the major cell types populating the CNS: neurons (NEUs), astrocytes (ASTs), microglia (MCGs), oligodendrocytes (OGDs), oligodendrocyte progenitors (OPCs), and endothelial cells (ENDs) [47-49]. The OB was the brain region with the highest number (37) of cell type markers differentially expressed between the different diets, followed by the BRN and BST, each with 12 DEGs, and the CBL with 9 DEGs (Figures 3D and S3B). Interestingly, most DEGs displaying enriched expression in animals fed the HFD or SCD were found in the OB, whereas most cell-type markers presenting enriched expression in NCD are the most abundant in the BST, followed by the BRN and CBL. Among all the cell type markers, the END marker Ftl1 is a notable exception, which consistently shows enriched expression in HFD or SCD compared to NCD across brain regions. Interestingly, increased brain expression levels of Ftl1 are associated with neurodegenerative diseases [50, 51].

Next, to gain deeper and unbiased insights into the diet-induced DEGs at the functional level, we performed GO over-representation analysis (see STAR Methods). We obtained between 6-57, 1-59, and 0-44 GO categories enriched for the HFD vs. NCD, SCD vs. NCD, and HFD vs. SCD, respectively (Data File S3). For the HFD vs. NCD and SCD vs. NCD comparisons, the GO-enriched categories related to the extracellular matrix and structure organization (OB, BRN, and BST), dynein binding (OB), calcium channel activity (OB), cognition (BRN), or endothelial development (BRN), are composed of genes enriched in animals fed an NCD. In contrast, GO-enriched categories composed of genes enriched in HFD-or SCD-fed animals are related to oxidative phosphorylation (OB, BRN, and CBL), ATP metabolism (OB, BRN, and CBL), mitochondrial activity and metabolism (OB, BRN, and CBL), or RNA-splicing mechanisms (CBL, and BST) (Figures 3E and F, Data File S3). For HFD vs. SCD comparison, we identified GO-enriched categories composed of genes enriched in animals fed an SCD, such as ones related to ribosomal processes in the OB, or RNA-splicing mechanisms in the BRN (Figure 3F, Data File S3). Finally, we also observed GO-enriched categories integrating genes enriched in animals fed an HFD, including several related to mitochondrial activity (OB).

Together, these data suggest that regardless of their macronutrient composition, short-term ingestion of semi-synthetic diets dysregulates a wide range of gene expression networks across the brain. Interestingly, many of the dysregulated pathways (e.g., extracellular matrix organization, oxidative phosphorylation, mitochondrial activity) are well-known to be associated with neuroinflammation in the brain, increasing the risk of developing neurodegenerative diseases [52].

### Effects of short-term consumption of semi-synthetic diets on weight gain, glucose tolerance, and adiposity

To further investigate the impact of short-term consumption of semi-synthetic diets, we assessed the body weight gain, glucose tolerance, and adiposity levels in mice fed the NCD, SCD, or HFD for 30 days in a different animal facility (see STAR Methods, Figure 4A). Mice fed the HFD gained significantly more body weight than those fed the NCD or the SCD (n=9-10 per group; two-way repeated measures ANOVA, Time: F(3,75) = 46.85, p < 0.0001; Diet: F(2,25) = 17.45, p < 0.0001; Interaction: F(6, 75)= 22.01, P<0.0001; Figures 4B and S4A). Next, we quantified the weights of the inguinal white adipose tissue (iWAT), gonadal WAT (gWAT), and interscapular brown adipose tissue (iBAT) at the end of the dietary intervention. While mice on the HFD showed higher amounts of iWAT and gWAT than those fed the NCD or the SCD, we observed no significant differences for the iBAT (n=8-9 per group; p<0.05, one-way ANOVA, BKY multiple-comparisons correction; Figure S4B). We then investigated the impact of the tested diets on glucose tolerance after 60 and 120 minutes of intraperitoneal bolus glucose injection. Only mice fed the HFD presented glucose intolerance (n=6 per group; two-way ANOVA, Time: F(4, 75) = 42.59, p < 0.0001; Diets: F(2, 75) = 15.57, p < 0.0001; interaction: F(8, 75) = 1.752, p = 0.1003, BKY multiple comparisons correction; Figure 4C), likely due to the interference of diet-derived circulating fats with the insulin receptor, which reduces glucose uptake by peripheral tissues [53, 54]. Together, these results show that only the short-term consumption of HFD negatively affects body weight, adiposity levels, and glucose tolerance.

**Fig 4.**
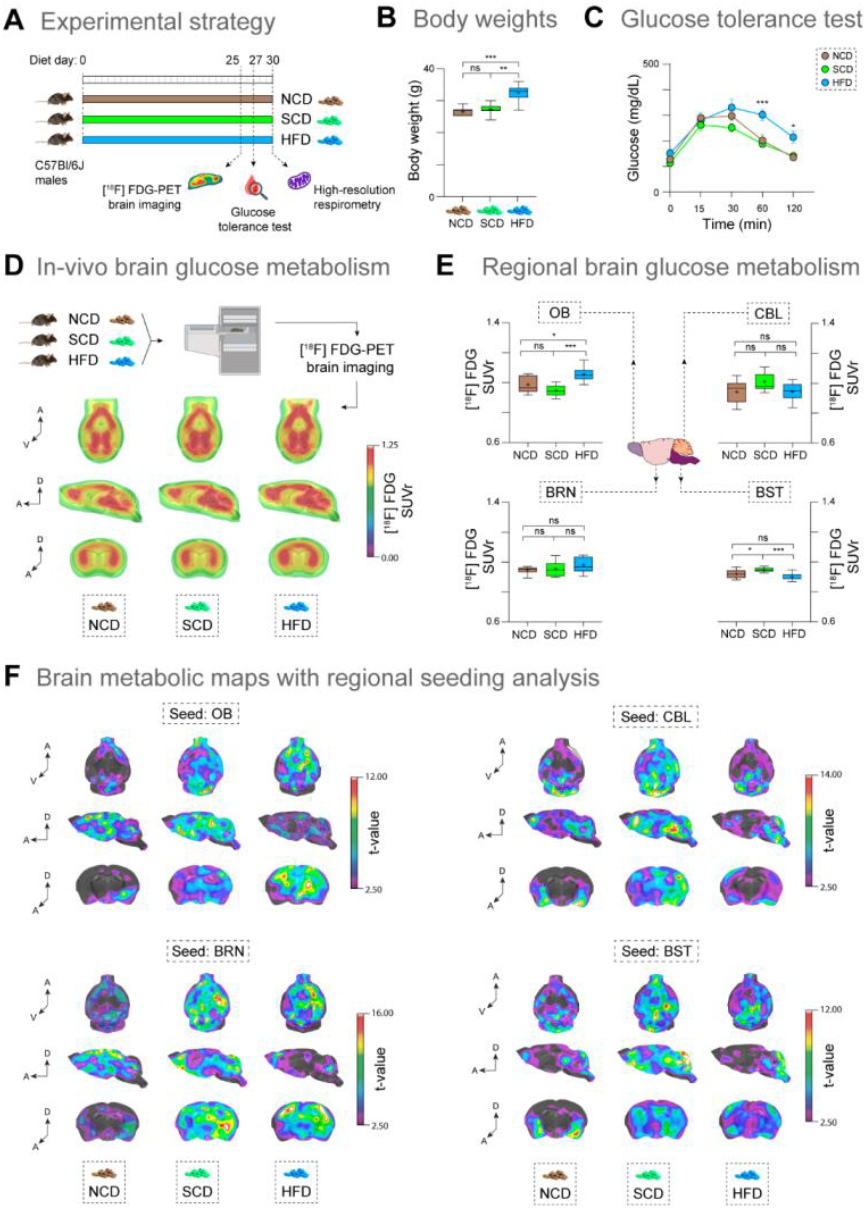
The synthetic-ingredient-based HFD induces metabolic adaptations in the brain. (A) Schematic view of the experimental design for the second part of our study. Mice were fed on different diets (NCD, SCD, HFD n=7-8 mice per group). In vivo brain glucose metabolism was assessed on day 25 of the dietary intervention, with [18F]-FDG-PET. The glucose tolerance test was performed on day 27. Brain mitochondrial function was analyzed in tissue homogenates on day 30 with high-resolution respirometry. (B) Comparison between the body weight of animals on a NCD, SCD, or HFD (n=7-8 per group). Asterisks indicate significant differences (one-way ANOVA, BKY multiple comparisons correction; ns, non-significant; **p<0.01; ***p<0.0001) (C) Glucose tolerance curves for animals on a NCD, SCD, and HFD right before glucose administration (t=0) and 15, 30, 60, and 120 min after an oral glucose administration. Animals fed the HFD have higher levels of glucose from minute 30 onwards, when compared with NCD and SCD. Asterisks indicate significant differences (two-way ANOVA, BKY multiple comparisons correction, n=6 per group): ns, non-significant; *p<0.05; ***p<0.0001). (D) Metabolic maps showing whole brain [18F]-FDG metabolism by SUVr of animals on a NCD, SCD and HFD. The pons region was used as reference. Warmer colors indicate higher SUVrs. (E) Specific templates for four regions of interest were used for separately quantify regional glucose metabolism in OB, CBL, BRN and BST. (F) The total brain glucose metabolism was used in a correlation analysis with a specific seed region to estimate glucose metabolism synchronicity between regions. Warmer colors indicate concomitant activation of brain areas and seeding point

### Short-term consumption of semi-synthetic diets affects brain metabolism and connectivity

Despite making only ∼2% of the body volume, the brain consumes ∼20% of the total glucose-derived energy [55]. This elevated energy consumption rate makes the brain highly sensitive to changes in dietary regimens [56]. Ex-vivo studies in rodents have shown that short- and long-term consumption of diets rich in fat or sucrose leads to brain metabolic changes [57-59]. However, whether the short-time consumption of semi-synthetic diets affects brain glucose metabolism remains largely unknown.

To answer these questions, we performed non-invasive brain glucose metabolism assessments with 18F-Fluorodeoxyglucose positron-emission tomography (FDG-PET) brain imaging after 25 days of dietary intervention (Figures 4A and 4D). Brain FDG-PET is widely used in clinics as an index of synaptic activity [60], and has an unprecedented translational value. We evaluated glucose metabolism in the same four cerebral regions (OB, BRN, CBL, and BST; n=9 per group) for which we performed RNA-seq in the section above. Our analysis revealed regional diet-induced changes in glucose metabolism in the OB and BST (p<0.05; one-way ANOVA, BKY multiple-comparisons correction; Figure 4E). In these two regions, glial cells are more abundant than neurons [61-63]. Interestingly, the OB of the mice on the HFD showed glucose hypermetabolism compared to mice fed the NCD or the SCD, and the BST of mice fed the SCD displayed hypermetabolism compared to animals on the NCD or the HFD (p<0.05; one-way ANOVA, BKY multiple-comparisons correction).

Next, to better understand the short-term dietary impact on brain metabolic connectivity, we used the OB, BRN, CBL, and BST as seed points for a network analysis (Figure 4F). The OB and the BRN seed point analyses showed hyperconnectivity in the animals fed the SCD and HFD, suggesting that these brain regions are consuming glucose in a more synchronized manner. This synchronicity could be due to short-term network adaptations to cope with increased energy substrate availability. Since glial cells outnumber neurons in these regions [61], one could argue that this hyper synchronicity is associated with this brain inflammatory state proposed above. The same analyses were conducted using brain regions enriched in neurons, such as the CBL, as the seed point. While the network exhibited by animals fed the NCD and HFD were similar, the animals fed the SCD displayed metabolic network hyper synchronicity. It is intriguing to consider that the metabolic network hyper synchronicity induced by HFD may depend on plastic changes occurring in glial cells [64-67]. Indeed, changes in brain metabolic connectivity due to astrocyte-specific modulation have been described before [64, 67].

Together, these results indicate that the short-term consumption of diets differing in macronutrient origin and composition can modulate regional brain glucose metabolism and metabolic networks, at least on a macro level. To our knowledge, this is the first in-vivo evidence of brain glucose hypermetabolism across many brain regions in such short dietary regimens using semi-synthetic diets, likely indicating a state of brain inflammation triggered by glial responses.

### Brain mitochondrial function is mildly affected by short-term dietary interventions

High-resolution respirometry was employed to assess mitochondrial performance in the OB, BRN, and the hypothalamus (HYP) of diet-treated mice, and several parameters associated with mitochondrial and non-mitochondrial respiration were measured (Figures 5A, and S5A). The respiratory acceptor control ratio (RCR, calculated by using the oxygen consumption in state3/state4) indicates the rate of mitochondria coupling and efficiency and may be altered under injury conditions [68]. We observed no significant differences in the RCR of the different brain regions between mice fed different diets (Figures 5B, and S5A). The uncoupling control ratio (UCR) is an index of the electron transfer system capacity related to the generation of heat by the mitochondria [69, 70]. The UCR was not significantly affected in animals on the SCD or HFD compared with NCD in the OB or the However, the UCR of the HYP was significantly higher in animals fed an NCD compared to animals on a SCD and HFD (one-way ANOVA, BKY multiple-comparisons correction; F(2, 15)=5.628, P=0.0150), consistent with the fact that this brain region is a critical regulator of energy metabolism in the brain, and it is highly affected by an overload of fat [6]. For the remaining four parameters linked to mitochondrial oxygen respiration (ATP-linked respiration, proton leak, and maximal respiration), we observed no significant differences in the RCR of the different brain regions between mice fed different diets. In the case of the non-mitochondrial oxygen consumption rate (OCR), we only observed significant differences (p<0.05; one-way ANOVA, BKY multiple-comparisons correction) for the HYP and OB, and only when comparing NCD against SCD or HFD. Processes related to this non-mitochondrial oxygen expenditure include the activity of oxidase enzymes, oxidation of odd-chain fatty acids, very long fatty acids, and amino acids [71, 72]. Under physiological conditions, this is a key process for the reoxidation of NADH, essential to the bioenergetic homeostasis in the cell [73].

**Fig 5.**
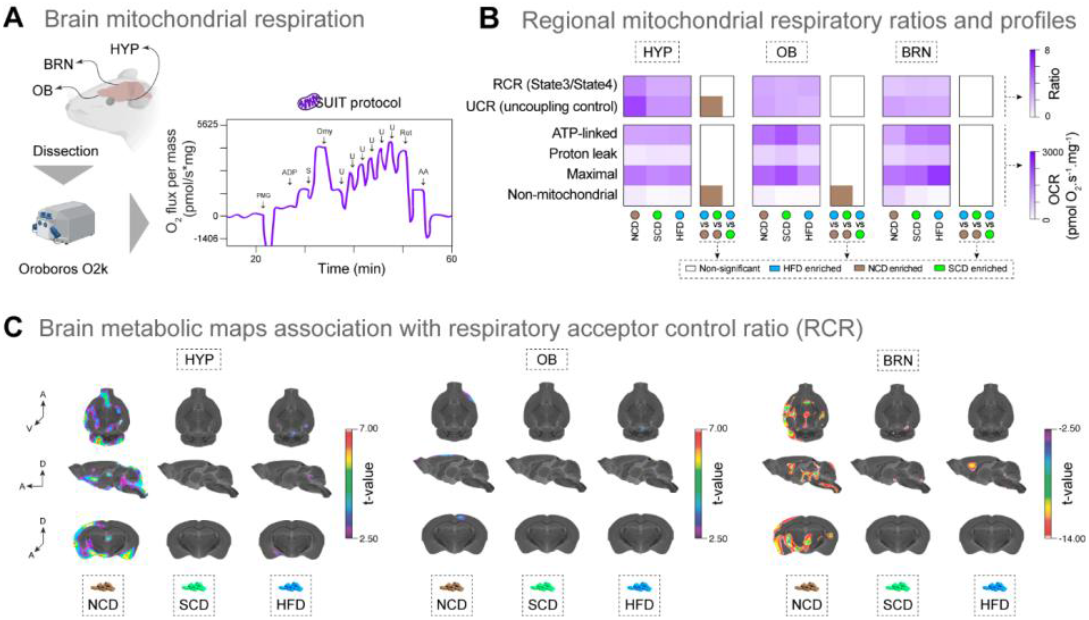
Brain mitochondrial function is mildly affected by the short-term consumption of synthetic-ingredient-based diets.

Schematic demonstration of brain regions used for high-resolution respirometry. To assess mitochondrial respiration in brain tissue homogenates, a substrate-inhibitor-titration (SUIT) protocol was used. Oxygen consumption at each stage of the protocol indicates specific mitochondrial properties. Mitochondria control ratios were calculated using the oxygen consumption in state3/state4 (RCR) and maximal respiration/leak (UCR). ATP-linked respiration is considered when all substrates, ADP and oxygen are available for a fully-coupled mitochondrial function. Proton leak measures are made in the presence of an ATP synthase inhibitor. Maximal respiration is achieved by titration with an electron transport system uncoupler. Non-mitochondrial is considered a residual oxygen consumption when the electron transport system and the ATP synthase are inhibited. (B) The HYP, OB, and BRN homogenates underwent testing with the SUIT protocol and had their mitochondrial parameters measured. Heatmap highlighting the mitochondrial functions measured. Similar shades of purple depict similar results.The box to the right of each heatmap indicates the statistical significance of inter-diet comparisons for each parameter (p<0.05; one-way ANOVA, BKY multiple comparisons correction, n = 4-7 per group). Comparisons with significantly higher values in NCD are indicated in brown. Non-significant responses are indicated in white. (C) Using in vivo glucose metabolism and the respiratory parameters, we estimated the brain glucose metabolism associations with respiratory acceptor control ratio in each brain region analyzed. Warmer colors indicate a positive correlation between RCR and brain glucose metabolism in the HYP and OB, and a negative correlation between RCR and brain glucose metabolism in the BRN.

Next, we conducted a voxel-wise association analysis integrating glucose metabolism and mitochondrial parameters. Interestingly, the hypothalamic mitochondrial parameters RCR and UCR are associated with hypothalamic (HYP) and cortical (BRN) glucose metabolism in the NCD group (Figures 5C, and S5C). In contrast, this association is not seen in SCD and HFD groups. This fact, specifically related to the hypothalamic region, can be a mechanism underlying the loss of homeostatic control the brain has on body weight, commonly seen in individuals consuming a HFD. The coupling between glucose metabolism and usage as an oxidative fuel seems abolished due to exposure to semi-synthetic SCD and HFD. No significant associations were found between glucose metabolism and the hypothalamic ROX (Figure S5D), suggesting that the hypothalamic non-mitochondrial usage of oxygen is not coupled to glucose metabolism. We conducted the same analysis with the OB mitochondrial parameters but found no associations between RCR or UCR and glucose metabolism (Figures 5C, and S5C). In contrast, when analyzing ROX and glucose metabolism, in the OB we found only small significant clusters (Figure S5D). These findings indicate that non-mitochondrial usage of oxygen in this region has little influence on glucose metabolism in the brain.

The whole-brain RCR was negatively associated with cerebral glucose metabolism in the NCD group but not in the others. This indicates that the FDG-PET signal is inversely associated with mitochondrial function, suggesting a glucose flux towards the glycolytic pathway independently of mitochondrial coupling. At the cellular level, this energetic flux is classically described in astrocytes, whereas in neurons we expect that lactate would be used for oxidative phosphorylation (and glucose for the pentose phosphate pathway) [74]. Corroborating our previous findings, this association is abolished by short-term SCD and HFD, which indicates an astrocytic plastic adaptation. Interestingly, the UCR in the BRN is negatively associated with glucose metabolism only in the HFD group, which indicates that regions consuming more glucose are likely less prone to deal with mitochondrial stress. This is not seen in the other groups. Regarding OX and glucose metabolism in the BRN, we found only small significant clusters.

Together, these results suggest that the usage of glucose, the main energetic substrate of the brain, is dependent on the nutrients available, and the cell type abundance in a given brain region. The inverse association between glucose metabolism and mitochondrial function suggests diet-induced adaptations are mostly driven by astrocytes, which are highly glycolytic cells.

## DISCUSSION

In mammals, previous studies investigating the effects of food on olfaction, brain metabolism, and overall health have predominantly focused on long-term dietary changes involving different macronutrient compositions that result in obesity [10, 21-25, 56, 59]. Here, we provide compelling evidence that short-term consumption of semi-synthetic diets, regardless of macronutrient composition and weight gain, can have profound effects on mouse olfactory physiology and odor-guided behaviors, transcriptional profiles of various cerebral regions, brain metabolism and connectivity, body weight, and glucose tolerance. Our results highlight that even short-term consumption of semi-synthetic foods can trigger early abnormalities in both the peripheral and central nervous systems, thus amplifying the susceptibility to neurological and other non-communicable diseases. Our study further substantiates the idea that the health effects of food are primarily dictated by its matrix, then its nutrient composition, with the quality of calories taking precedence over their quantity [75].

Overall, mice fed the semi-synthetic diets showed substantially altered olfactory preferences and risk assessment episodes, suggesting that the effects of diet on olfactory sensory and cognitive functions may manifest more rapidly than previously thought. These strong diet-induced effects on odor-guided behaviors occur even in the absence of obesity, in line with a previous study of long-term exposure to a 45% high-fat diet [24]. Interestingly, obese humans also display altered olfactory preferences and overall lower olfactory sensitivity and function [76, 77], but whether the same remains true in non-obese individuals with poor diet or poor metabolic health is vastly unknown. Our results also provide intriguing insights into the potential mechanisms underlying these dietary effects on behavior. The observed impairments in odorant detection (measured by the EOG responses) and the associated behavioral deficits could be attributed to changes in the peripheral detection of odorants in the neuroepithelium of the mouse WOM, possibly mediated by alterations in the response of OSNs to odorants to diet-derived metabolic internal signals [10, 20, 22, 41]. Indeed, our EOGS findings are consistent with the impairment of the peripheral detection of odorants in the mouse nose after the long-term consumption of unhealthy high-fat or high-fructose diets (Thiebaud, Johnson et al. 2014, Riviere, Soubeyre et al. 2016). Furthermore, the transcriptional responses associated with apoptosis and inflammatory responses in the mouse nose following consumption of semi-synthetic diets suggest that these diets may induce physiological dysregulation of odor perception, possibly through diet-induced dysregulation of neuroregenerative, inflammatory, or apoptotic processes [23, 25, 42-44].

Internal metabolic signals are also powerful regulators of many physiological processes throughout the brain, including key areas involved in the perception of smell and related behaviors, such as the OB, the piriform cortex, and the hypothalamus [10, 20, 41, 78-80]. Our findings align with this, as the electro-olfactogram responses of OSNs to the odorants accounted for about half of the variability in these behaviors, suggesting the involvement of additional modulatory mechanisms in the brain. In this context, the consumption of different macronutrients in the diet is known to modulate gene expression patterns, thereby affecting brain physiology, and significantly impacting metabolic health and brain function [37, 45, 46]. In our transcriptomic analyses, we identified DEGs in the WOM and across various brain regions in response to diet consumption, suggesting that the impact of diet on brain function is organ and region-specific. The OB, a key brain region involved in regulating olfactory responses [10, 19], registered the highest number of DEGs across all the comparisons (including the WOM), yielding ∼53-fold higher number of DEGs for the comparison between the two semi-synthetic diets, thus suggesting that this brain region may be particularly sensitive to dietary interventions and be a key modulator of the olfactory responses. The GO over-representation analysis further revealed that the short-term consumption of semi-synthetic diets dysregulates gene expression networks across the brain, including those related to energy metabolism, oxidative phosphorylation, RNA-splicing, myelination, calcium-channel activity, and others. These findings are consistent with the idea that different diets have distinct effects on different cell types and brain regions, leading to diverse functional outcomes [49, 65, 81-84].

Past studies in rodents indicate that long-term exposure to HFD or binge access to a high-sucrose diet can significantly impact brain glucose metabolism and provoke cerebral inflammation, even in the absence of obesity [36, 56, 58, 59, 65, 83]. In our experiments, the short-term consumption of HFD has profound effects on body weight, glucose tolerance, and adiposity levels. In alignment with these findings, our study revealed that short-term consumption of semi-synthetic diets can modulate regional brain glucose metabolism and metabolic networks, and in the absence of obesity. This was evidenced by the hypermetabolism observed in the OB and BST of mice fed the HFD and SCD, respectively. Brain metabolism can react to peripheral cues to adapt its responses depending on external stimuli. Our results indicate that glial cells of OB had their main markers altered by HFD and SCD, as a consequence of diet-induced metabolic alterations. Specifically, the astrocytic marker Slc1a2 [85], which encodes GLT-1 main trigger for glucose uptake in astrocytes, was altered under the different dietary regimens, which may be directly related to increased glucose metabolism in OB. Moreover, the lack of changes in glucose metabolism in regions where neurons outnumber glial cells (such as the CBL), combined with the fact that glial cells are ultra-plastic in handling glucose [86], suggest that the observed in-vivo glucose hypermetabolism in such a short regimen indicates glial adaptive responses. Indeed, recent studies have shown that glial cells contribute to the FDG-PET signal [64, 66, 67, 87]. Thus, such dynamic responses would likely have as their cellular source the glial cells, particularly astrocytes – the most abundant glial cell in the gray matter. These findings suggest that diet can rapidly influence brain function, potentially through mechanisms involving glial cells, which are known to be highly plastic and play a critical role in brain metabolism [88]. Additionally, evidence from clinical studies indicates that early glucose hypermetabolism seen in neurodegenerative disorders is associated with reactive astrogliosis and microglial activation [87, 89], suggesting that even a short-term consumption of semi-synthetic diets may cause brain inflammatory changes.

Obesity and metabolic syndrome, caused by the long-term consumption of unhealthy diets, can lead to mitochondrial dysfunction in the brain [90, 91]. Our findings indicate that short-term dietary interventions can subtly influence brain mitochondrial function, aligning with the early stages of this pathophysiological process. While the respiratory acceptor control ratio and uncoupling control ratio were not significantly affected in the four broad brain regions, significant changes were observed in the hypothalamus. This suggests that short-term consumption of semi-synthetic diets can influence mitochondrial function in specific brain regions, potentially contributing to the observed changes in brain metabolism and connectivity.

While our study provides novel insights into the impact of short-term semi-synthetic diet consumption on olfactory function and gene expression profiles in various brain regions, there three major several limitations to consider. First, our study was conducted in mice, and while they are a widely used model for human diseases, the results may not fully translate to humans (or other species) due to differences in physiology and metabolism. Second, we focused on three specific diets (NCD, SCD, and HFD), leaving the effects of other types of diets or variations in macronutrient composition unexplored. Third, our study was of short duration, and the medium-or long-term effects of these diets on olfactory function and brain gene expression could differ.

In conclusion, our findings highlight the intricate and multifaceted influence of diet on sensory perception, behavior, brain function, and metabolic health. They suggest that diet can swiftly alter brain function, potentially through mechanisms involving the highly adaptable glial cells, which are pivotal in brain metabolism. This underscores the complex interplay between diet and brain function, emphasizing the need for further research to fully understand these relationships. Ultimately, these findings underscore the importance of developing dietary interventions that can enhance metabolic health and optimize brain function, further emphasizing the profound impact of diet on overall health and well-being.

Our findings open several avenues for future research. It would be beneficial to examine the long-term effects of these diets and to investigate the impact of other diet types or variations in macronutrient composition on olfactory function and brain gene expression. Our findings also suggest that, the olfactory system could serve as a potential early indicator or sentinel for the onset of metabolic disorders, is in addition to its recognized role as a biomarker for aging, neurodegenerative issues, and respiratory diseases [92, 93]. Additionally, more comprehensive studies are needed to fully understand the cellular and molecular mechanisms underlying the observed changes in olfactory function and brain gene expression. This could involve more in-depth investigations into the role of different cell types and other elements of the brain’s gene expression network. Ultimately, a deeper understanding of the impact of diet on olfactory function and brain gene expression could guide the development of dietary interventions to promote health and prevent disease.

## MATERIAL AND METHODS

### Animals

The animals used in the assays for this study were male C57Bl/6J mice (The Jackson Laboratory, Stock # 00664) maintained in group-housed conditions on a 12:12 h light:dark schedule (lights on at 07.00 hours). The use and care of animals used in this study was approved by the Internal Animal Care and Use Committee (IACUC) of Monell Chemical Senses Center, and by the IACUC of the Universidade Federal do Rio Grande do Sul (UFRGS).

### Dietary interventions

In this study, we used three diets differing in macronutrient composition (Figure 1A, Data File S1): a grain-based normal chow diet (NCD, 54% carbohydrate, 32% protein, 14% fat; #8604 Teklad Rodent diet - Envigo), a semi-synthetic control diet (SCD, 70% carbohydrate, 20% protein, 10% fat; #D12450B - Research Diets Inc.), and a semi-synthetic high-fat diet (HFD, 20% carbohydrate, 20% protein, 60% fat; #D12492 - Research Diets Inc.). Mice had ad libitum access to food and water throughout the duration of the study. Upon arrival at the Monell Chemical Senses Center animal facility, male C57Bl/6J mice (5 weeks old) were habituated for 5 days, and then randomly assigned to one of the three dietary experimental groups: NCD (n=85), SCD (n=72), and HFD (n=79). The dietary intervention lasted 23-58 days (see Figure 1). Upon arrival at the UFRGS animal facility, C57BL/6J male mice (10-12 weeks old) were randomly assigned to one of the three dietary experimental groups: NCD (n=9), SCD (n=10), and HFD (n=9).

### Odorants

The odorants used in this study were purchased from Sigma-Aldrich at the highest purity available. Their names are shown below, followed by their abbreviations in parentheses: (+)-menthone (+MNT), ethyl butyrate (EBT), hexanethiol (HXT), hexanol (HXO), indole (IND), isoamylamine (IAA), linalool (LIN), pentyl acetate (PAC), trimethylamine (TMA), vanillin (VAN). For the behavioral assays, odorants were either diluted in water to a concentration of 85 mM, or first dissolved in dimethyl sulfoxide (DMSO) to make a 1 M stock solution and then diluted in water to a concentration of 85 mM, immediately before the assays. For the EOG recordings, odorants were first dissolved in DMSO to make a 5 M stock solution, then diluted in water to obtain odorant solutions at 10^−2^ M in 5 ml final volumes in sealed 50 ml glass bottles. As a control (0 M odorant), a solution of DMSO equivalent to the concentration in the 10^−1^ M odorant solution was used.

### Olfactory preference test and behavioral scoring

The olfactory preference tests were performed at Monell Chemical Senses Center, 22-32 days after the start of the dietary treatment on 9-10 week-old mice, as previously described [39] and with minor modifications (see below). Each mouse was assayed only once to avoid possible bias due to learning, and data for each odorant consisted of 7-12 mice. On the day of testing, mice were brought to the experimental room 120-180 minutes prior to the onset of the dark cycle (at 19:00) and habituated for 5–10 min. Mice were then separated into new cages (one mouse per cage) without food or water and habituated again for 15-30 minutes. Exposure to odorants was done by gently introducing a stainless-steel mesh tea ball containing a cotton ball impregnated with 50 μL of double-distilled water (H2O) or odorant (85 mM in H2O, equivalent to 4.25 μmol), into one end of the cage. Videos were recorded from the top, for slightly over 3 minutes, using a GoPro HERO6 Black camera (Waterproof Digital Action Camera for Travel with Touch Screen 4K HD Video). All behavioral assays were conducted during the last 2 hours of the light cycle, between 17:00 and 19:00. Prior to scoring, all videos were cropped with a custom-made tool developed in our lab – the “Video Crop” tool (https://github.com/NeethuVenugopal/videocrop) – to eliminate areas around the mouse cage that will interfere with the downstream automated video tracking done using Ethovision XT software (version 11, Noldus Information Technology). Briefly, the “Video Crop” tool is a script encoding a video cropping tool that provides a graphical user interface (GUI) for users to select a specific region of interest (ROI) in a video. By interactively clicking and dragging the mouse, users can define the ROI on the video frames. Once the selection is confirmed, the script crops the frames within the ROI and saves the cropped video as an output file. This tool enables users to extract specific portions of a video, focusing only on the desired content. The behavioral parameters of each mouse/video were scored by a non-experimenter blinded to the odor used in the assay. Three behavioral parameters were scored during a 3-minute period, starting from introducing the odorant-containing metal tea ball:

- Olfactory investigation time (OIT) represents the cumulative duration of olfactory investigation during the assay duration. This parameter is a surrogate measure for valence [40], and scored using the Ethovision XT software (version 11, Noldus Information Technology) with the video tracking done using the nose-point of the mice. An automated detection setting based on greyscale analysis was used to detect mice in this analysis, and olfactory investigation was considered only if the nose-point of the mice was touching the odorant-containing tea ball.
- Risk assessment (RAS) represents the number of episodes the mouse displays the flat-back/stretch-attend response, followed by a sniff in the direction of the stimulus, during the assay duration. This parameter is a surrogate measure for stress [40], and was scored manually.
- Velocity (VEL) represents the mean velocity at which the mouse walked/ran during the assay duration. This parameter is a surrogate measure for activity [40], and scored using the Ethovision XT software (version 11, Noldus Information Technology) with the video tracking done using the center-point of the mouse. An automated detection setting based on greyscale analysis was used to detect mice in this analysis (refer to Ethovision guide).

### Electro-olfactogram recordings

The electro-olfactogram (EOG) recordings were performed at Monell Chemical Senses Center, 38-47 days after the start of the dietary treatment on 10-11 week-old mice, as previously described [94] and with minor modifications (see below). Data for each dietary group consisted of 7-8 mice. The mice were euthanized with CO2 and decapitated, and their heads were sagitally bisected at the center of the nasal septum. The septum was removed to expose the olfactory turbinates in the nasal cavity. The bisected heads were quickly transferred to a recording setup, where a stream of humidified air flowed (3 l/min) over the tissue. In these experiments, we used the odorants PAC, VAN, IND, HXT, IAA, +MNT, LIN, EBT, and TMA. The headspace from each odorant solution was injected with a Picospritzer (Parker Hannifin, Cleveland, OH, USA) into the air stream flowing over the OE to stimulate OSNs. EOG recordings were performed in the order of PAC, VAN, IND, HXT, IAA, PAC, +MNT, H2O, LIN, EBT, TMA, DMSO control solution, and PAC again as a control for recording stability over time. To record the EOGs, two electrodes were placed on the surfaces of turbinate II and IIB respectively using either the left or right half of the head at similar positions in all mice [95]. The signals were recorded with two DP-301 amplifiers (Warner Instruments, Hamden, CT, USA), and the 1 kHz low-pass-filtered signal was digitized at 2 kHz with a Micro1401 mkII digitizer and Signal version 5.01 software (Cambridge Electronic Design, Milton, Cambridge, England). Recordings were analyzed by averaging across the four channels recorded from both sides of the nasal cavity. If there was more than a 25% reduction in the response amplitude between the first PAC and the last PAC recording from a given electrode, the data were excluded. On occasions, mice had nasal cavities with strongly deviating septa, which typically gave small responses, and these data were also excluded.

### Tissue dissection, RNA extraction, and RNA-sequencing

Tissue dissections were performed at Monell Chemical Senses Center, 52-58 days after the start of the dietary treatment on 12-13 week-old mice. All animals were sacrificed by cervical dislocation between 14:00 and 16:00). From each animal, the following organs were collected and processed for mRNA-seq: whole olfactory mucosa (WOM, dissected as described in [96]), olfactory bulb (OB), brain (BRN, which includes the telencephalon and diencephalon), cerebellum (CBL), brainstem (BST, which consists of the mesencephalon, pons, and myelencephalon). The organs were immediately frozen and kept at -80°C until further processing. WOM, OB, BRN, CBL, and BST samples were homogenized using a tissue homogenizer (OMNI International) in Qiazol (Qiagen). Total RNA for the WOM, OB, BRN, CBL, and BST samples was extracted using the Lipid RNeasy Lipid Tissue Mini Kit (Qiagen), according to the manufacturer’s protocol. mRNA was prepared for sequencing using the TruSeq stranded mRNA sample preparation kit (Illumina), with a selected insert size of 120–210 bp. Stranded libraries were sequenced on an Illumina HiSeq 4000, generating paired-end 150 bp sequencing reads, with an average depth of 45 ± 1.4 (SEM) million reads.

### Short read alignment to the reference genome and transcriptome

The quality of the reads was assessed using FastQC (KBase). Sample Fastq files were aligned to the mouse reference genome GRCm38.p6 (GCA_000001635.8) using TopHat2 (version 2.1.1) [97] with 2 mismatches allowed. Reads were retained only if uniquely mapped to the genome. We used HTSeq-count (0.9.1, -t exon and –m union) to obtain the number of reads that were mapped to each gene in Gencode M24. Bigwig files were generated from bam files for visualization using bam2wig.py function RSeQC v3.0.1 [98].

### Differential expression analysis and functional annotation

Sample transcriptome data quality was assessed by evaluating homogeneity and similarity between samples. We used variance stabilizing transformation (vst) and regularized log transformation(rlog) functions (DESeq2 package, v1.26.0) [99] to transform the raw count data, (dist) function to calculate sample-to -sample distances. We plotted heatmaps of distance matrix along with PCA of 1000 HVG and the union of the top 1000 most expressed genes to identify putative outlier samples. These samples if present were excluded from the following differential expression analyses. We also used DESeq2 (v1.26.0) to normalize the raw count matrix according to sequencing depth and RNA composition (median of ratios method), then to perform differential expression analysis. Differential expression was tested using the DESeq function from the DESeq2 library, which employs the Wald test to determine if the log2 fold change in gene expression across two groups of samples is higher than expected by chance. Finally, DESeq does p-value correction (adjusted p-value) for multiple testing. Genes with a padj < 0.05 and |log2FC |> 1 were considered as differentially expressed. Gene over-representation analysis was performed using ClusterProfiler package (v4.0.0) (PMID: 34557778). We used Gene Ontology enrichment (enrichGO) and Simplify functions, with ont = “BP”, “MF” or “CC” and pAdjustMethod = “BH” parameters to assess which Biological Processes, Molecular Functions, or Cellular Components were affected in our lists of differentially expressed genes.

### ^18^F-Fluorodeoxyglucose micro positron-emission tomography (FDG-PET) scanning

These experiments were performed at UFRGS, 25 days after the start of the dietary treatment on 14-16 week-old mice, as previously described [67] and with minor modifications (see below). After overnight fasting (12 hours, from 08:00 PM to 08:00 AM), the mice received an intravenous injection (0.4 mL) of FDG-PET (mean ± s.d.: 0.26 ± 0.006 mCi) in the tail vein. Then, each mouse was returned to its home cage for a 40-minute period of conscious (awake) in-vivo uptake of FDG, which was followed by a 10 minute static acquisition under anesthesia (2 % isoflurane at 0.5 L/min oxygen flow). FDG-PET measurements were performed on a Triumph™ micro-PET [LabPET-4, TriFoil Imaging, Northridge, CA, USA (for LabPET-4 technical information see [100]). The brain was positioned in the center of the field of view (FOV; 3.75 cm), and the body temperature was maintained at 36.5 ± 1 °C. All data was reconstructed using the maximum likelihood estimation method (MLEM-3D) algorithm with 20 iterations. Each FDG-PET image was reconstructed with a voxel size of 0.2 × 0.2 × 0.2 mm and spatially normalized into an FDG-PET template using brain normalization in PMOD v3.8 and the Fuse It Tool (PFUSEIT) (PMOD Technologies, Zurich, Switzerland). The following imaging analysis was conducted using the minc-tools software (www.bic.mni.mcgill.ca/ServicesSoftware/MINC). The standardized uptake value ratio (SUVr) was calculated using pons as the reference region. Mean SUVrs of the four analyzed brain regions were extracted using a predefined volume of interest (VOI) template.

### Glucose tolerance test

The glucose tolerance test experiments were performed at UFRGS, 27 days after the start of the dietary treatment on 11-week-old mice, as previously described [101], and with minor modifications (see below). Mice were orally administered with a 2 g/kg glucose solution after an overnight fast (12 hours, from 08:00 PM to 08:00 AM). Tail vein blood glucose was measured with the Accu-Chek Active meter over 2 hours (0, 15, 30, 60, and 120 min).

### High-resolution mitochondrial respirometry

These experiments were performed at UFRGS 30 days after the start of the dietary treatment on 13-15-week-old mice, as previously described [102], and with minor modifications (see below). Mice were euthanized by decapitation, had their HYP, OB and BRN rapidly dissected and put on ice. Representative slices of all regions (in the case of cortex, herein called “whole brain”) or the whole tissue (in the case of hypothalamus and olfactory bulb) were weighted, homogenized 10%(w/v) in respiration buffer (Mir05 - 110 mM sucrose, 60 mM potassium lactobionate, 20 mM taurine, 10 mM monobasic potassium phosphate, 3 mM magnesium chloride, 20 mM HEPES, 1 mM EGTA, and 0.1% (w/v) BSA at pH 7.1) using a glass potter (10-12 strokes on ice). Samples were transferred to an oxygraph-2k (O2k, Oroboros Instruments, Innsbruck, Austria) in a final concentration of 100-200 μg total protein per chamber at 37 °C and initial O2 concentration of around 200 μM. Substrate, uncoupler, inhibitor titration (SUIT) protocol was carried out in Mir05 (adapted from [103]): respiratory states were induced with pyruvate 5 mM, malate 4 mM, and glutamate 10 mM. Oxidative phosphorylation activity was measured with ADP 500 μM and succinate 10 mM. Oligomycin 0.2 μg/mL (Omy), an ATP synthase inhibitor, was used to determine leak respiration. Titration of the uncoupler carbonyl cyanide m-chlorophenyl hydrazone (CCCP) 0.5 μM allowed verification of maximal O2 consumption. Complex I inhibition was obtained with rotenone 0.5 μM (R). Residual consumption was induced with antimycin A 2.5 μM (AA). The remaining oxygen consumption after R and AA was considered non-mitochondrial respiration. Respirometry was analyzed with the DatLab software (v6.1.0.7. Oroboros Instruments). Respiratory acceptor control ratio (RCR) was defined by the coupling state of mitochondria obtained by the division of the O2 flux in the presence of substrates and ADP (OXPHOS – state 3) by the O2 flux after Omy addition (state 4). OXPHOS capacity is the oxidation coupled to phosphorylation in the presence of saturating O2, ADP, and substrate concentration. Proton leak is a dissipative component of respiration that is not available for performing biochemical work and is thus related to heat production. Uncoupling control ratio (UCR) is the ratio between the maximum respiratory capacity achieved under uncoupler titration (CCCP) and the O2 flux during proton leak [104].

### Statistical Analysis

Statistical analyses were done using R Statistical Software (version 4.1.0.), GraphPad Prism (version 8.0.0), and Origin software version 8.5 (Origin Lab, Northampton, MA, USA), or PAlaeontological STatistics (version 4.06, http://folk.uio.no/ohammer/past/). For the principal component analysis (Fig. 2C and S2A,B), the data matrix was standardized, and correlation matrixes used to compute the eigenvalues and eigenvectors. Results were analyzed by one-way or two-way ANOVA, and P-values were computed with Benjamini, Krieger and Yekutieli (BKY) multiple comparisons correction (P-values <0.05 were considered significant.).

## DATA AVAILABILITY

RNA-seq raw data (Fastq and raw counts files) have been deposited and are publicly available as of the peer-reviewed publication date at GEO under the study accession number GSE218162. All original code and scripts for the Video Crop” tool have been deposited on Github and can be found at https://github.com/NeethuVenugopal/videocrop. All original code and scripts for the RNA-seq and the brain imaging analyses is available on demand.

## ACKNOWLEDGMENTS

We would like to acknowledge Dr. Amber L. Alhadeff, Dr. Darren W. Logan, Prof. Kunio Kondoh, Dr. Julie Mennella, and the members of the Saraiva Lab and Zimmer Lab for their insightful inputs on data analysis and/or comments on the manuscript. We are thankful to Susie Huang for preliminary bioinformatic analysis, and the Integrated Genomics Services of Sidra Medicine for performing their assistance with the RNA-sequencing.

## AUTHOR CONTRIBUTIONS

M.M., D.G.S., S.K., and B.B. performed experiments, analyzed data, and contributed to the writing of the initial version of the manuscript. B.B., H.L., and AK. performed experiments. A.A-N., R.H., G.T.V., J.C. da C., S.S.Y.H., N.V., and D.M. analyzed data. J.R., and M.T. analyzed data, and helped write the manuscript. E.Z. and L.R.S. conceived and supervised the project, analyzed data, and wrote the initial and final versions of the manuscript.

## REFERENCES

1. Popkin, B. M. Am J Clin Nutr 84, 289–298 (2006).

2. Wiley, A. S. in Basics in Human Evolution (ed Michael P. Muehlenbein) 393–404 (Academic Press, 2015).

3. McCrickerd, K. & Forde, C. G. Obes Rev 17, 18–29 (2016).

4. Boesveldt, S. & Parma, V. Cell Tissue Res 383, 559–567 (2021).

5. Hall, K. D. et al. Cell Metab 30, 67–77 e63 (2019).

6. Loos, R. J. F. & Yeo, G. S. H. Nat Rev Genet 23, 120–133 (2022).

7. Kopp, W. Diabetes Metab Syndr Obes 12, 2221–2236 (2019).

8. de Araujo, T. P. et al. Int J Environ Res Public Health 18 (2021).

9. Goncalves, N. et al. JAMA Neurol 80, 142–150 (2023).

10. Palouzier-Paulignan, B. et al. Chem Senses 37, 769–797 (2012).

11. Boesveldt, S. & de Graaf, K. Perception 46, 307–319 (2017).

12. Zoon, H. F. et al. Foods 5 (2016).

13. Soria-Gomez, E. et al. Nat Neurosci 17, 407–415 (2014).

14. Herz, R. S. Brain Sci 6 (2016).

15. Kontaris, I. et al. Front Behav Neurosci 14, 35 (2020).

16. Kershaw, J. C. & Mattes, R. D. World J Otorhinolaryngol Head Neck Surg 4, 3–10 (2018).

17. Kong, I. G. et al. PLoS One 11, e0164495 (2016).

18. Rawal, S. et al. Nutrients 13 (2021).

19. Guzman-Ruiz, M. A. et al. Cell Mol Neurobiol (2021).

20. Jovanovic, P. & Riera, C. E. Trends Endocrinol Metab 33, 281–291 (2022).

21. Lietzau, G. et al. ACS Chem Neurosci 11, 3590–3602 (2020).

22. Riviere, S. et al. Sci Rep 6, 34011 (2016).

23. Chelette, B. M. et al. J Physiol 600, 1473–1495 (2022).

24. Takase, K. et al. Obesity (Silver Spring) 24, 886–894 (2016).

25. Thiebaud, N. et al. J Neurosci 34, 6970–6984 (2014).

26. Cootes, T. A. et al. Cell Rep 41, 111638 (2022).

27. Tremblay, A. J. et al. Am J Clin Nutr 98, 32–41 (2013).

28. Kuipers, E. N. et al. Am J Physiol Endocrinol Metab 317, E820–E830 (2019).

29. Wiedemann, M. S. et al. Am J Physiol Endocrinol Metab 305, E388–395 (2013).

30. Edwards, L. M. et al. FASEB J 25, 1088–1096 (2011).

31. Clara, R. et al. J Cell Physiol 232, 167–175 (2017).

32. Zeng, T. et al. Biogerontology 20, 837–848 (2019).

33. Shang, Y. et al. Lipids 52, 499–511 (2017).

34. Eudave, D. M. et al. Neurobiol Stress 9, 1–8 (2018).

35. Arnold, S. E. et al. Neurobiol Dis 67, 79–87 (2014).

36. Kothari, V. et al. Biochim Biophys Acta Mol Basis Dis 1863, 499–508 (2017).

37. Sarangi, M. & Dus, M. Front Behav Neurosci 15, 746299 (2021).

38. Parma, V. et al. Chem Senses 45, 609–622 (2020).

39. Saraiva, L. R. et al. Proc Natl Acad Sci U S A 113, E3300–3306 (2016).

40. Manoel, D. et al. Curr Biol 31, 2809–2818 e2803 (2021).

41. Bryche, B. et al. Cell Tissue Res 384, 589–605 (2021).

42. Ueha, R. et al. Front Aging Neurosci 10, 86 (2018).

43. Chen, M. et al. Proc Natl Acad Sci U S A 114, 8089–8094 (2017).

44. Robinson, A. M. et al. Laryngoscope 112, 1431–1435 (2002).

45. Vaziri, A. & Dus, M. Neurochem Int 149, 105099 (2021).

46. Ravi, S. et al. J Nutr 145, 841–846 (2015).

47. De Jager, P. L. et al. Nat Neurosci 17, 1156–1163 (2014).

48. McKenzie, A. T. et al. Sci Rep 8, 8868 (2018).

49. Bondareva, O. et al. Nat Metab 4, 1591–1610 (2022).

50. David, S. et al. Antioxid Redox Signal 37, 150–170 (2022).

51. Kim, Y. K. et al. Int J Mol Sci 24 (2022).

52. Picca, A. et al. Antioxidants (Basel) 9 (2020).

53. Shulman, G. I. J Clin Invest 106, 171–176 (2000).

54. Sears, B. & Perry, M. Lipids Health Dis 14, 121 (2015).

55. Magistretti, P. J. & Allaman, I. Neuron 86, 883–901 (2015).

56. Iozzo, P. & Guzzardi, M. A. Endocr Connect 8, R169–R183 (2019).

57. Jais, A. et al. Cell 166, 1338–1340 (2016).

58. Estadella, D. et al. Lipids Health Dis 10, 168 (2011).

59. Patkar, O. L. et al. Sci Rep 11, 11252 (2021).

60. Stoessl, A. J. Nat Neurosci 20, 382–384 (2017).

61. Herculano-Houzel, S. Glia 62, 1377–1391 (2014).

62. Bandeira, F. et al. Proc Natl Acad Sci U S A 106, 14108–14113 (2009).

63. von Bartheld, C. S. et al. J Comp Neurol 524, 3865–3895 (2016).

64. Zimmer, E. R. et al. Nat Neurosci 20, 393–395 (2017).

65. Garcia-Caceres, C. et al. Cell 166, 867–880 (2016).

66. Zimmer, E. R. et al. Sci Transl Med 14, eabm8302 (2022).

67. Rocha, A. et al. Eur J Nucl Med Mol Imaging 49, 2251–2264 (2022).

68. Gilmer, L. K. et al. J Neurotrauma 27, 939–950 (2010).

69. Demine, S. et al. Cells 8 (2019).

70. Gnaiger, E. Bioenerg Commun 2020 (2020).

71. Chacko, B. K. et al. Clin Sci (Lond) 127, 367–373 (2014).

72. Banh, R. S. et al. Nat Cell Biol 18, 803–813 (2016).

73. Herst, P. M. et al. Biochim Biophys Acta 1656, 79–87 (2004).

74. Pellerin, L. & Magistretti, P. J. J Cereb Blood Flow Metab 32, 1152–1166 (2012).

75. Fardet, A. & Rock, E. Eur J Nutr 61, 2239–2253 (2022).

76. Stafford, L. D. & Whittle, A. Chem Senses 40, 279–284 (2015).

77. Velluzzi, F. et al. Nutrients 14 (2022).

78. Al Koborssy, D. et al. Brain Struct Funct 224, 315–336 (2019).

79. Nogi, Y. et al. Sci Rep 10, 890 (2020).

80. Horio, N. & Liberles, S. D. Nature 592, 262–266 (2021).

81. Duking, T. et al. Sci Adv 8, eabo7639 (2022).

82. Garcia-Caceres, C. et al. Nat Neurosci 22, 7–14 (2019).

83. Jais, A. et al. Cell 165, 882–895 (2016).

84. Munji, R. N. et al. Nat Neurosci 22, 1892–1902 (2019).

85. Perego, C. et al. J Neurochem 75, 1076–1084 (2000).

86. Magistretti, P. J. J Exp Biol 209, 2304–2311 (2006).

87. Xiang, X. et al. Sci Transl Med 13, eabe5640 (2021).

88. Jha, M. K. & Morrison, B. M. Exp Neurol 309, 23–31 (2018).

89. Salvado, G. et al. Eur J Nucl Med Mol Imaging 49, 4567–4579 (2022).

90. Cavaliere, G. et al. Front Cell Neurosci 13, 509 (2019).

91. Schmitt, L. O. & Gaspar, J. M. Metabolites 13 (2023).

92. Dan, X. et al. Ageing Res Rev 70, 101416 (2021).

93. Gerkin, R. C. et al. Chem Senses 46 (2021).

94. Cygnar, K. D. et al. J Vis Exp (2010).

95. Barrios, A. W. et al. Front Neuroanat 8, 63 (2014).

96. Ruiz Tejada Segura, M. L. et al. Cell Rep 38, 110547 (2022).

97. Kim, D. et al. Genome Biol 14, R36 (2013).

98. Wang, L. et al. Bioinformatics 28, 2184–2185 (2012).

99. Love, M. I. et al. Genome Biol 15, 550 (2014).

100. Bergeron, M. et al. Phys Med Biol 59, 661–678 (2014).

101. Souza, C. G. et al. Food Funct 4, 1271–1276 (2013).

102. da Silva, J. S. et al. Mol Neurobiol 57, 4790–4809 (2020).

103. Pesta, D. & Gnaiger, E. Methods Mol Biol 810, 25–58 (2012).

104. Brand, M. D. & Nicholls, D. G. Biochem J 435, 297–312 (2011).

