## Supplementary Information for "Short-term consumption of ultra-processed semi-synthetic diets impairs the sense of smell and brain metabolism in mice"

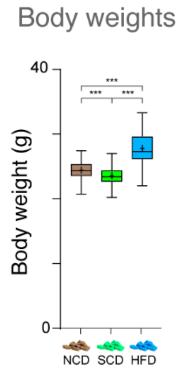


**Figure S1. Effects of short-term consumption of diets on body weight. Related to Figure 1 and Data File S1.** The body weights differ for all comparisons, with animals fed a SCD displaying the lower weight, followed by NCD, and HFD (n=78-84 per group). Asterisks indicate significant differences (one-way ANOVA, BKY multiple comparisons correction; ***p<0.0001).

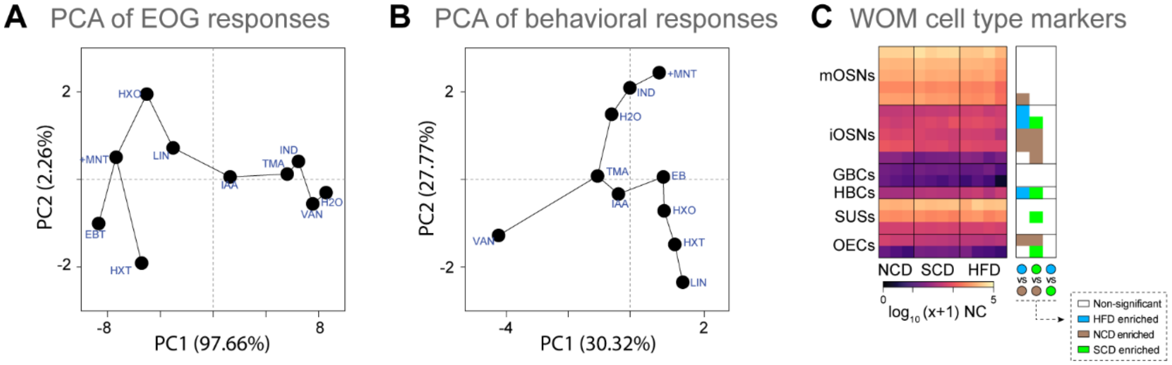


**Figure S2. Ultra-processed diets affect odorant detection and induce transcriptional dysregulations in the mouse nose. Related to Figure 2 and Data File S2.** (A,B) Principal component analysis (PCA) was performed on the data matrixes from the EOG responses (A) and the odor-guided behaviors (B) for the nine tested odorants and water. Percentages of the variance explained by the PCs are indicated in parentheses. (C) Heatmap (left panel) showing the normalized gene expression levels of WOM cell type marker genes. Right panel: a graphical display in the form of a heatmap summarizing the statistical significance of inter-diet comparisons. DEGs (p-adj<0.05) enriched in mice fed on HFD, NCD, or SCD are represented in blue, brown, or green, respectively. mOSNs, mature olfactory sensory neurons (OSNs); iOSNs, immature OSNs; GBCs, globose basal cells; HBCs, horizontal basal cells; SUSs, sustentacular cells; OECs, olfactory ensheathing cells.

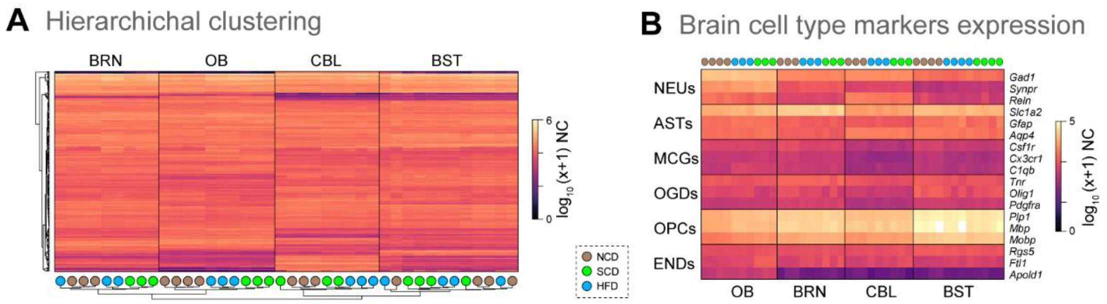


**Figure S3. Ultra-processed diets differentially affect the transcriptome of various brain regions. Related to Figure 3 and Data File S3.** (A) Hierarchical clustering on the expression values of the union of the top 1000 most expressed genes across all brain regions samples. (B) Heatmap showing the normalized expression levels of genes that are known markers for six brain cell types: NEUs, neurons; ASTs, astrocytes; MCGs, microglial cells; OGDs, oligodendrocytes; OPCs, oligodendrocyte progenitor cells; ENDs, endothelial cells.

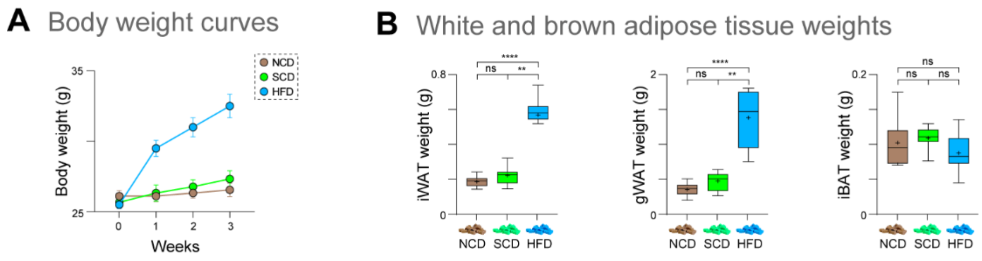


**Figure S4. High-fat diet regimen induces metabolic adaptations. Related to Figure 4 and Data File S4.** (A) The body weight did not differ between NCD and SCD, but it was significantly higher in HFD animals since week 1 compared to NCD and SCD (n=9-10 per group; two-way repeated measures ANOVA, BKY multiple comparisons correction). (B) After four weeks of diet, inguinal white adipose tissue (iWAT), gonadal white adipose tissue (gWAT), and interscapular brown adipose tissue (iBAT) weights were measured. iWAT and gWAT weights were increased in HFD-fed mice compared to SCD and NCD, while no changes in iBAT weight were observed (n=8-9 per group). Asterisks indicate significant differences (one-way ANOVA, BKY multiple comparisons correction; ns, non-significant; **p<0.01; ***p<0.0001).

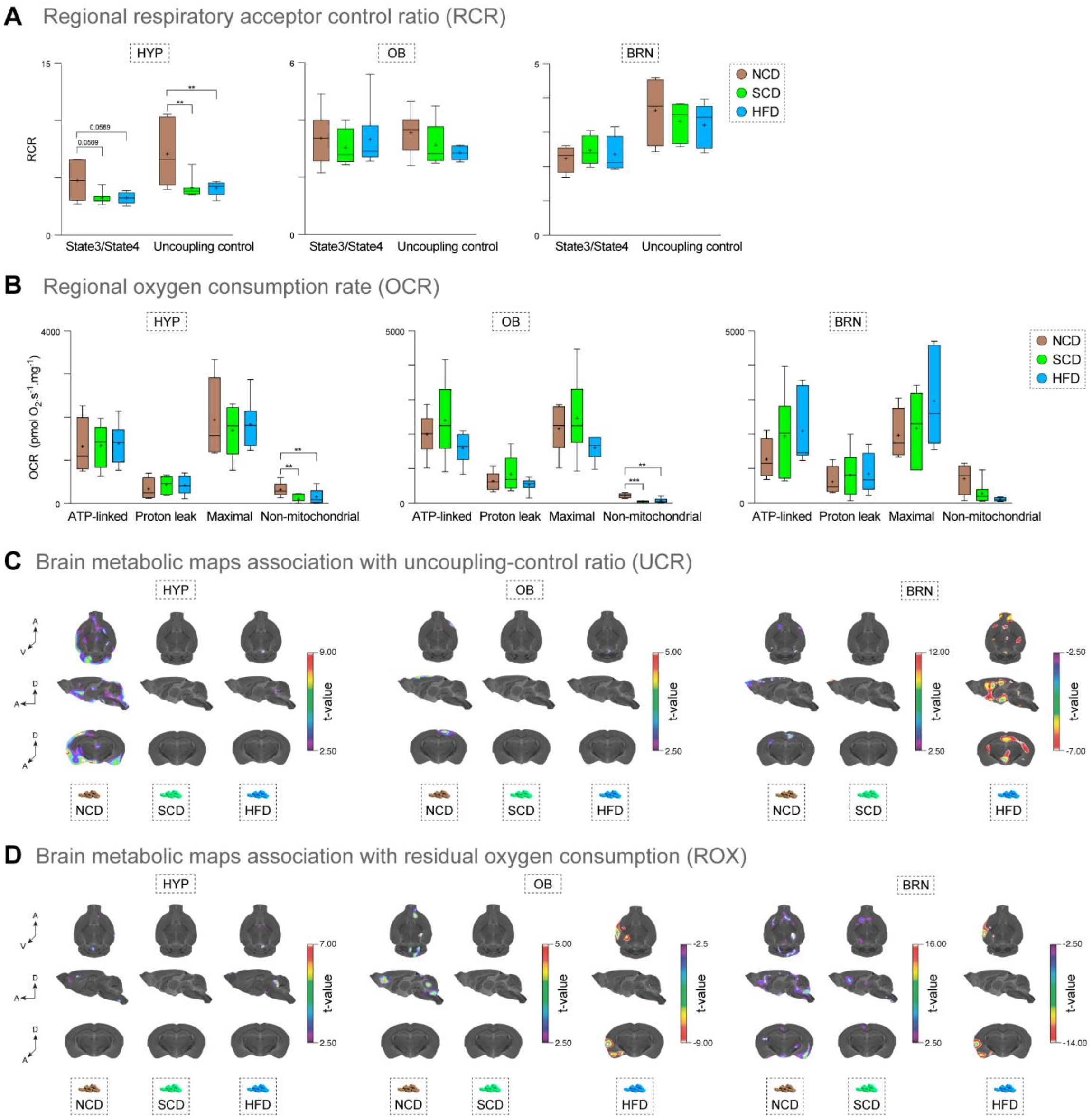


**Figure S5. Brain mitochondrial function is mildly affected by the short-term consumption of a high-fat diet. Related to Figure 5 and Data File S5.** (A) The HYP, OB, and BRN homogenates were tested with a SUIT protocol. Oxygen consumption at each stage of the protocol indicates specific mitochondrial properties. Mitochondria control ratios were calculated using the oxygen consumption in state3/state4 (RCR) and maximal respiration/leak (UCR). The boxplots (n=4-7 per group) shown here are related to the heatmap from Figure 5B. (B) ATP-linked respiration is considered when all substrates, ADP and oxygen are available for a fully-coupled mitochondrial function. Proton leak measures are made in the presence of an ATP-synthase inhibitor. Maximal respiration is achieved by titration with an electron transport system uncoupler. Non-mitochondrial is considered a residual oxygen consumption when the electron transport system and the ATP synthase are inhibited. The boxplots (n=4-7 per group) shown here are related to the heatmap from Figure 5B. (C) Using in vivo glucose metabolism and the respiratory parameters, we estimated the brain glucose metabolism associations with UCR in each brain region analyzed. Warmer colors indicate a positive correlation between UCR and brain glucose metabolism in the HYP and OB for all diets, and for BRN under NCD and SCD diets. For HFD, there is a negative correlation between UCR and brain glucose metabolism in the BRN. (D) Brain glucose metabolism associations with regional residual oxygen consumption (ROX) showed a sparse signal in HYP, OB and BRN showed a positive association in NCS and SCD groups, but a negative association in HDF group, in regions indicated with warmer colors. OCR: oxygen consumption rate.
